# Simulation of Ca^2+^ oscillations in astrocytes mediated by amyloid beta in Alzheimer’s disease

**DOI:** 10.1101/2020.03.18.996843

**Authors:** Huayi Gao, Langzhou Liu, Shangbin Chen

## Abstract

Disruptions of astrocyte Ca^2+^ signaling is important in Alzheimer’s disease (AD) with the unclear mechanism of amyloid beta peptide (Aβ). We have modified our previous computational model of spontaneous Ca^2+^ oscillations in astrocytes to investigate the effects of Aβ on intracellular Ca^2+^ dynamics. The simulation results have shown consistence with the previous experiments. Aβ can increase the resting concentration of intracellular Ca^2+^ and change the regime of Ca^2+^ oscillations by activating L-type voltage-gated calcium channels and the metabolic glutamate receptors, or by increasing ryanodine receptors sensitivity and Ca^2+^ leakage, respectively. This work have provided a toolkit to study the influence of Aβ on intracellular Ca^2+^ dynamics in AD. It is helpful for understanding the toxic role of Aβ during the progression of AD.

**Statement of Significance:** Alzheimer’s disease (AD) is the most common neurodegenerative disease with the unclear mechanism of amyloid beta peptide (Aβ). This work have implemented a computational model to address the Ca^2+^ dynamics of astrocyte mediated by Aβ with the four different pathways: voltage-gated calcium channels, metabotropic glutamate receptors 5, ryanodine receptor channels and membrane leak. The Ca^2+^ oscillations and bifurcation diagram indicate that astrocytes exhibit ionic excitability mediated by Aβ and become the potential targets of Aβ neurotoxicity. We expect this shared computational model would advance the understanding of AD.

## Introduction

Alzheimer’s disease (AD) is the most common neurodegenerative disease which could result in cognitive decline and memory loss [1]. Amyloid beta peptide (Aβ) deposit and its neurotoxicity associated with AD. In particular, Aβ is involved in the disruption of Ca^2+^ regulation in astrocytes. Astrocytes are hypothesized as the primary target of Aβ neurotoxicity [2]. Recently, a computational model has been applied to investigate the effects of Aβ on Ca^2+^ regulation in an AD environment [3]. So far, there has been no model addressing the unique features of astrocyte. Based on our previous model [4] and the literature reported data [5-8], we have implemented the relatively comprehensive simulations on Ca^2+^ oscillations in astrocytes mediated by Aβ.

## Methods on the simulation

In the model astrocyte (see Figure 1), different types of voltage-gated calcium channels (VGCCs) [5] form the Ca^2+^ influx *J*_V*GCC*_ from the extracellular space (ECS) to the intracellular space (ICS). The electrophysiological properties of these VGCCs were described by the Hodgkin-Huxley (HH) equations. Only the L-type VGCC current may be medicated by Aβ [6]. To investigate the effects of Aβ, we use parameter *α* to represent a fixed level of Aβ concentration presenting in the environment. This parameter is also used in the following equations. In addition, we use *A*_*VL*_ to control the strength of Aβ effects on the pathway of L-type VGCC current, so the total Ca^2+^ current is recorded as:

**FIGURE 1.**
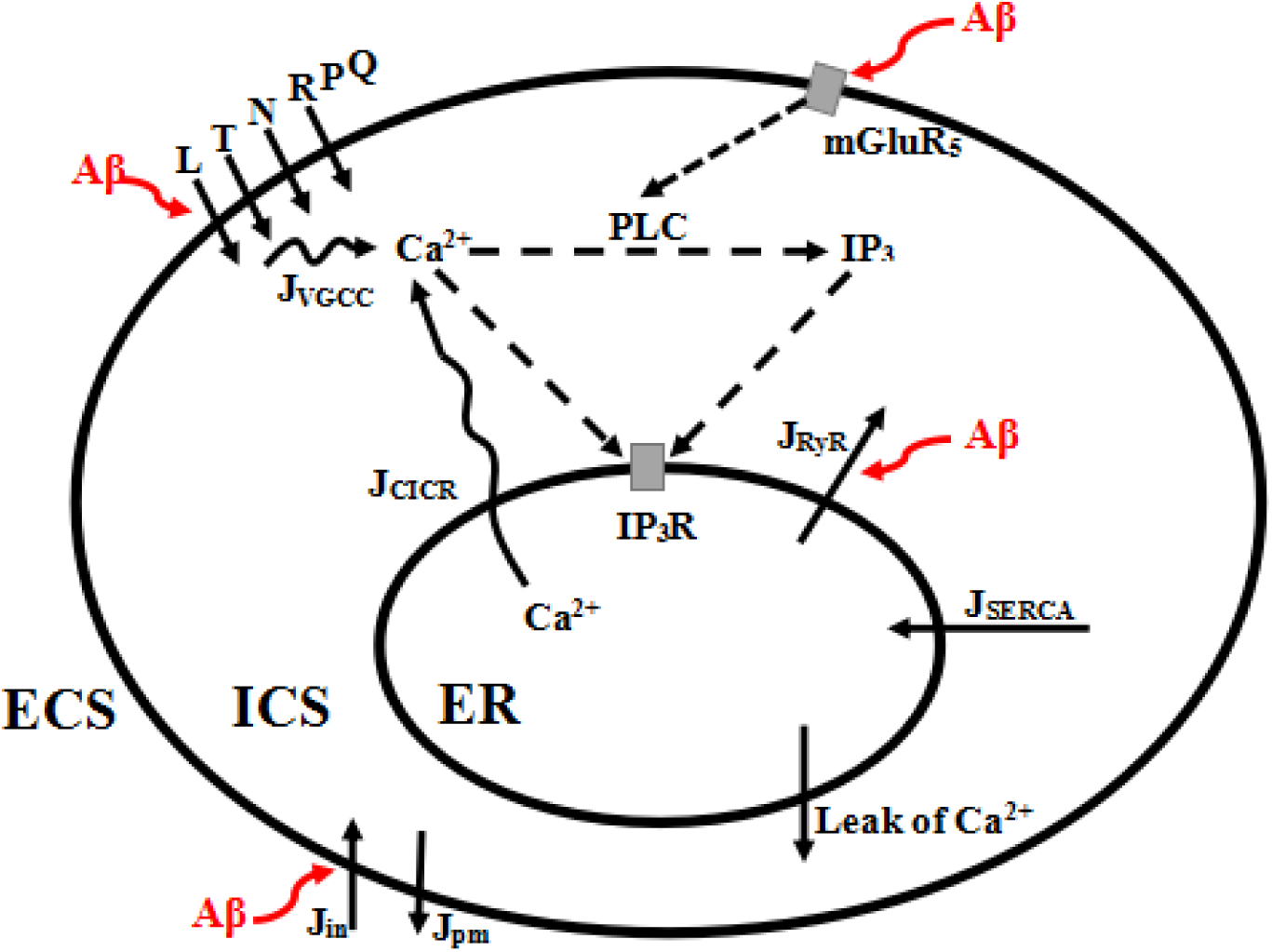
The model astrocyte with different Ca^2+^ fluxes. Aβ may mediate the L-type voltage-gated calcium channels (VGCC), metabotropic glutamate receptors 5 (mGluR_5_), ryanodine receptor (RyR) channels and membrane leak *J*_*in*_. In this model, cytoplasmic Ca^2+^ enhances the production of inositol 1,4,5-triphosphate (IP_3_) catalyzed by phospholipase C (PLC). Cytoplasmic Ca^2+^ and IP_3_ mediate IP_3_ receptors (IP_3_R), inducingCa^2+^ flow out of the endoplasmic reticulum (ER) by calcium-induced calcium release (CICR). *J*_*SERCA*_ represents ER Ca^2+^ filling by the sarco-endoplasmic reticulum Ca^2+^ ATPase (SERCA). The ‘‘leak’’ arrow from ER indicates the leak flux due to the concentration gradient.

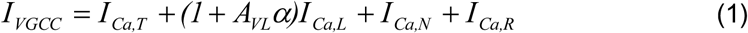

And the corresponding flux is:

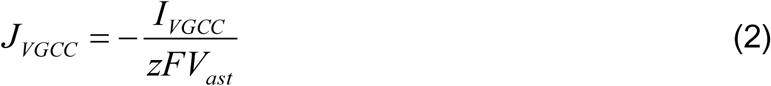

Referred to the previous work [3, 4], the other Ca^2+^ fluxes *J*_*CICR*_, *J*_*SERCA*_, *J*_*RyR*_, *J*_*LER*_, *J*_*in*_ and *J*_*pm*_ are written as:

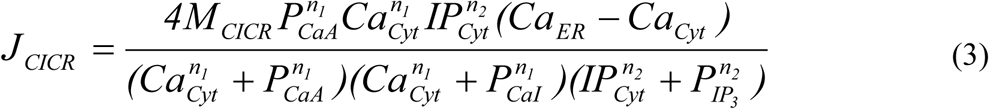

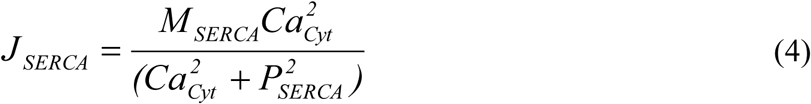

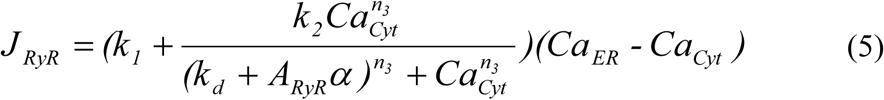

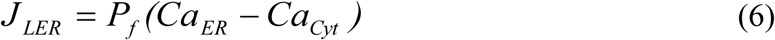

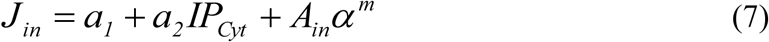

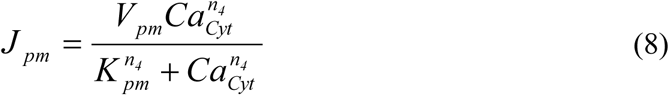

Where *J*_*LER*_ is a leak from the ER to the cytoplasm. In *J*_*in*,_ *J*_*RyR*_ and *J*_*PLC*_, we also use *α* to represent the effects of Aβ, and use *A*_*in*_, *A*_*RyR*_, *A*_*mGluR*_ to control the strength of these different effects. The relevant parameters are listed and briefly introduced in Table 1.

**TABLE 1.** Variables and Parameters Used in the Model.

| Symbol | Equation | Description | Value or unit |
| --- | --- | --- | --- |
| $I$ | Eq. 1 | Ionic current | fA |
| $A_{VL}$ | Eq. 1 | The strength of the effect of Aβ on the L-type current in $I_{VGCC}$ | 1 |
| $\alpha$ | Eq. 1 | Fixed level of Aβ concentration present in the environment | [ 0 , 1 ) |
| $J_{VGCC}$ | Eq. 2 | Influx of extracellular Ca<sup>2+</sup> into cytosol via VGCCs | 0 μM/s (t = 0) |
| $z$ | Eq. 2 | Valence of Ca<sup>2+</sup> ion | 2 |
| $F$ | Eq. 2 | Faraday's constant | 96,485 Coul/mole |
| $V_{ast}$ | Eq. 2 | Volume of an astrocyte | $3.49 \times 10^{-13}$ l |
| $J_{CICR}$ | Eq. 3 | IP<sub>3</sub>-mediated CICR flux to the cytosol from the ER | 0 μM/s (t = 0) |
| $M_{CICR}$ | Eq. 3 | Maximum flux of calcium ions into the cytosol | 10 s<sup>-1</sup> |
| $P_{CaA}$ | Eq. 3 | Activating affinity | 0.15 μM |
| $n_1$ | Eq. 3 | Hill coefficient | 2.02 |
| $n_2$ | Eq. 3 | Hill coefficient | 2.2 |
| $Ca_{Cyt}$ | Eq. 3 | Ca<sup>2+</sup> concentration in cytosol | 0.1 μM (t = 0) |
| $IP_{Cyt}$ | Eq. 3 | IP<sub>3</sub> concentration in cytosol | 0.1 μM (t = 0) |
| $Ca_{ER}$ | Eq. 3 | Ca<sup>2+</sup> concentration in ER | 1.5 μM (t = 0) |
| $P_{Cal}$ | Eq.3 | Inhibiting affinity | 0.15 μM |
| $P_{IP3}$ | Eq. 3 | Half-saturation constant for IP<sub>3</sub> activation of IP<sub>3</sub>R | 0.1 μM |
| $J_{SERCA}$ | Eq. 4 | The filling with Ca<sup>2+</sup> from the cytosol to ER | 0 μM/s (t = 0) |
| $M_{SERCA}$ | Eq. 4 | Maximum flux across SERCA | 15.0 μM/s |
| $P_{SERCA}$ | Eq. 4 | Half-saturation constant for SERCA activation | 0.1 μM |
| $J_{RyR}$ | Eq. 5 | RyR flux to the cytosol from the ER | 0 μM/s (t = 0) |
| $k_1$ | Eq. 5 | Zero calcium concentration level leak | 0.013 s<sup>-1</sup> |
| $k_2$ | Eq. 5 | Maximal rate of the channel | 0.18 s<sup>-1</sup> |
| $k_d$ | Eq. 5 | RyR channel sensitivity for CICR | 0.13 μM |
| $n_3$ | Eq. 5 | Hill coefficient | 3 |
| $A_{RyR}$ | Eq. 5 | The strength of the effect of Aβ on J<sub>RyR</sub> | 1 μM |
| $J_{LER}$ | Eq. 6 | Leak flux from the ER into the cytosol | 0 μM/s (t = 0) |
| $P_f$ | Eq. 6 | Rate of leak flux from the ER into the cytosol | 0.01 s<sup>-1</sup> |
| $J_{in}$ | Eq. 7 | Membrane leak flux into the cytosol | 0 μM/s (t = 0) |
| $a_1$ | Eq. 7 | Parameter | 0.003 μM/s |
| $a_2$ | Eq. 7 | Parameter | 0.02 s<sup>-1</sup> |
| $A_{in}$ | Eq. 7 | The strength of the effect of Aβ on J<sub>in</sub> | 1 μM/s |
| $m$ | Eq. 7 | Cooperativity coefficient | 3.5 |
| $J_{pm}$ | Eq. 8 | Flux of $\text{Ca}^{2+}$ via plasma membrane pump | 0 $\mu\text{M/s}$ ($t = 0$) |
| $V_{pm}$ | Eq. 8 | Maximal velocity | 4.2 $\mu\text{M/s}$ |
| $K_{pm}$ | Eq. 8 | Channel sensitivity | 0.425 $\mu\text{M}$ |
| $n_4$ | Eq. 8 | Hill coefficient | 2 |
| $J_{PLC}$ | Eq. 9 | Production of IP3 | 0 $\mu\text{M/s}$ ($t = 0$) |
| $A_{mGluR}$ | Eq. 9 | The strength of the effect of $\text{A}\beta$ on $\text{mGluR}_5$ | 1 |
| $M_{PLC}$ | Eq. 9 | Maximum production rate of PLC | 0.05 $\mu\text{M/s}$ |
| $P_{PCa}$ | Eq. 9 | Half-saturation constant for calcium activation of PLC | 0.3 $\mu\text{M}$ |
| $P_{deg}$ | Eq. 12 | Rate of IP3 degradation | 0.08 $\text{s}^{-1}$ |

And the IP_3_ production rate *J*_*PLC*_ is determined as:

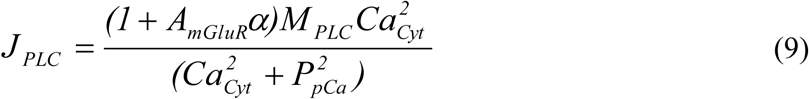

So the Ca^2+^ concentration in the cytosol (*Ca*_*Cyt*_), Ca^2+^ concentration in the ER (*Ca*_*ER*_) and the IP_3_ concentration in the cell (*IP*_*Cyt*_) are described by the ordinary differential equations:

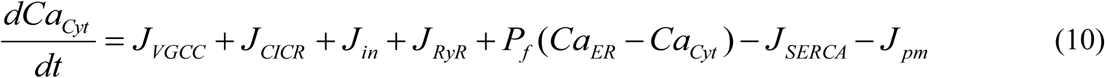

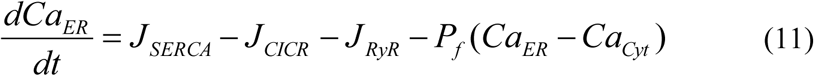

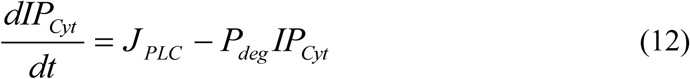

The MATLAB codes (MATLAB R2017a, The MathWorks, Natick, MA) are prepared to solve the ordinary differential equations and shared in the Supporting Material.

## Results and Discussion

Our model can reproduce typical Ca^2+^ oscillations with or without the influence of Aβ in astrocytes (see the Figure 2). By setting different parameters of Aβ in the Eqs. 1, 5, 7 and 9, we can check the dynamics of *Ca*_*Cyt*_, *Ca*_*ER*_ and *IP*_*Cyt*_. For simplicity, we only focus on the change of *Ca*_*Cyt*_. Although, the four different pathways (i.e. VGCC, mGluR5, RyR and membrane leak *J*_*in*_) mediated by Aβ act to affect astrocytes with different properties, the most significant common features are three points: increasing the frequency of Ca^2+^ oscillations, lowering the membrane threshold for Ca^2+^ oscillations and increasing the stable state concentration of Ca^2+^.

**FIGURE 2.**
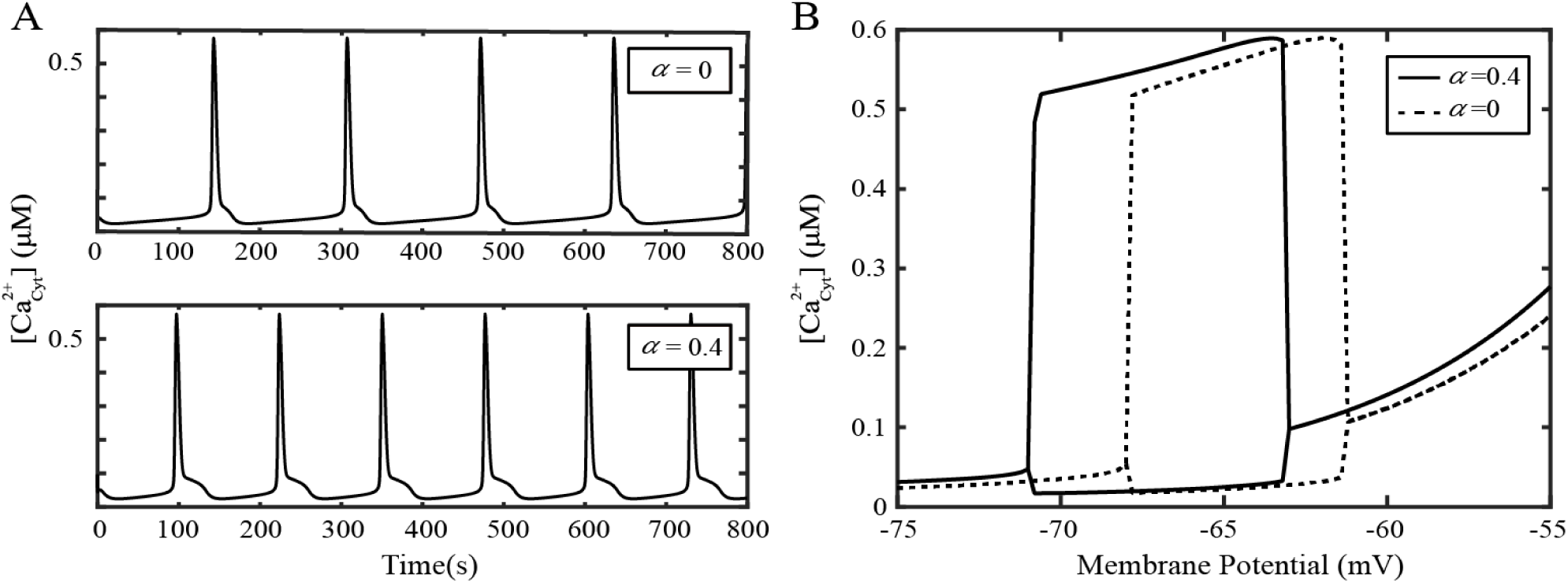
Comparative study with or without Aβ effects on Ca^2+^ oscillations in astrocyte (*α* = 0 means without Aβ). (A) The frequency of Ca^2+^ oscillations is increased by Aβ at the membrane potential of −65 mV. (B) The bifurcation diagram shows that Aβ can alter the membrane potential regime of Ca^2+^ oscillations and increase the stable state concentration of Ca^2+^.

AD astrocytes associated with senile plaques demonstrate Ca^2+^ hyperactivity in the form of aberrant Ca^2+^ oscillations [7]. The simulated results are well consistent with the reported experiments. Aβ mainly enlarges the membrane potential range and increases the resting Ca^2+^ at low steady state by increasing *J*_*in*_. According to the reference [8], the clustering of mGluR_5_ caused by Aβ lead to the shift of the oscillating range to lower potential. The increasing amplitude of the Ca^2+^ oscillations is majorly caused by the increasing sensitivity of RyR channel. Aβ increases the resting Ca^2+^ at the high steady state and shift the oscillating range to lower potential by activating L-type VGCC. Aβ can activate the L-type channels and increase the concentration of intracellular Ca^2+^. In turn, the block of L-type VGCC can protect cells from the detrimental effects of Aβ. Astrocytes exhibit ionic excitability [5] mediated by Aβ. This work indicates that preventing the effects from Aβ can prevent the development of Alzheimer’s disease.

Although some previous experiments indicated that Aβ had no effect on intracellular calcium in neurons but caused striking changes in adjacent astrocytes [2], the major limitation of this work is ignoring the effect of Aβ on neurons [9]. This is could be studied in the future. This is a generic model on the cellular mechanisms of Aβ on Ca^2+^ regulation in AD. The integration of the four different pathways mediated by Aβ has improved the biological plausibility of AD simulation. This shared computational model and the reproducible research would truly advance the understanding of AD [10].

## Author Contributions

S.C. and H.G. designed the research; H.G. and S.C. performed the research; H.G. S.C. and L.L. wrote the paper.

## Acknowledgements

This work was supported by the Science Fund for Creative Research Groups (Grant No. 61721092), the National Natural Science Foundation of China (Grant Nos. 61371014, 91749209), and the Director Fund of WNLO.

